# Ecological consequences of intraspecific variation in coevolutionary systems

**DOI:** 10.1101/2020.03.09.984161

**Authors:** Athmanathan Senthilnathan, Sergey Gavrilets

## Abstract

The patterns and outcomes of coevolution are expected to depend on intraspecific trait variation. Various evolutionary factors can change this variation in time. As a result, modeling coevolutionary processes solely in terms of mean trait values may not be sufficient; one may need to study the dynamics of the whole trait distribution. Here, we develop a theoretical framework for studying the effects of evolving intraspecific variation in two-species coevolutionary systems. In particular, we build and study mathematical models of competition, exploiter-victim interactions, and mutualism in which the strength of within- and between-species interactions depends on the difference in continuously varying traits. We use analytical approximations based on the invasion analysis and supplement it with a numerical method. We find that intraspecific variation can be maintained if stabilizing selection is weak in at least one species. When intraspecific variation is maintained, stable coexistence is promoted by small ranges of interspecific interaction in two-species competition and mutualism, and large ranges in exploiter-victim interactions. We show that trait distributions can become multimodal. Our approach and results contribute to the understanding of the ecological consequences of intraspecific variation in coevolutionary systems by exploring its effects on population densities and trait distributions.

## Introduction

Coevolution, that is, reciprocal evolutionary changes in ecologically interacting species, is a major research area where ecology and evolution come together (Futuyma and Slatkin, 1983; Nuismer, 2017; Thompson, 1994). Although coevolution is difficult to demonstrate in natural systems (Janzen, 1980; Nuismer *et al.*, 2010), its importance in nature is well established (Brouat *et al.*, 2001; Clayton *et al.*, 1999; Davies and Brooke, 1988; Soler *et al.*, 2001; Toju and Sota, 2006). Coevolution explains a number of evolutionary phenomena including character displacement in competing species, evolutionary arms race between exploiters and victims, and trait correlation between mutualist partners (Anderson and Johnson, 2008; Benkman *et al.*, 2003; Pfennig and Pfennig, 2009). The geographic mosaic theory of coevolution (GMTC) further highlights the relevance of coevolution by incorporating spatial variation in natural selection and strength of ecological interactions (Thompson, 2005). Through GMTC, coevolution can explain patterns of clinal variation, local adaptation, and stability of mutualisms in the presence of costs (Gavrilets and Michalakis, 2008; Gómez *et al.*, 2009; Gomulkiewicz *et al.*, 2000; Nuismer *et al.*, 2000).

Coevolution is not only a very important process but is also complex due to the intricate relationship between evolutionary and ecological processes. Because of its complexity, there is a strong need for a mathematical theory capturing this relationship. Correspondingly, various population genetics, quantitative genetics, and adaptive dynamics models and methods have been used to study eco-evolutionary dynamics in specific two-species systems: competition (Kremer and Klausmeier, 2017; Leimar *et al.*, 2013; Roughgarden, 1976; Slatkin, 1980; Taper and Case, 1992), mutualism (Akçay, 2016; Ferrière *et al.*, 2002, 2007; Foster and Kokko, 2006), and exploiter-victim interactions (Abrams, 2000; Doebeli, 1997; Fleischer *et al.*, 2018; Gavrilets, 1997; Gavrilets and Michalakis, 2008; Nuismer *et al.*, 2005; Schreiber *et al.*, 2016). Doebeli and Dieckmann (2000); Kopp and Gavrilets (2006); Yoder and Nuismer (2010) modeled all three types of two-species interactions within the same framework. For example, using a weak selection approximation and numerical simulations, Kopp and Gavrilets (2006) studied the dynamics of allele frequencies, means, and variances of a trait controlled by several diploid diallelic loci. Yoder and Nuismer (2010) modeled how trait variation in a metapopulation changes due to coevolution using a quantitative genetics approach and an individual-based model. Doebeli and Dieckmann (2000) used the adaptive dynamics approach to study phenotypic diversification due to ecological interactions. They also verified the analytic predictions using individual-based models. These and other similar mathematical models explicated the effects of the eco-evolutionary feedback in coevolution.

Both biological intuition and mathematical models tell us that the time-scales, patterns, and outcomes of coevolution should depend on within-species genetic and phenotypic variation. (Albert *et al.*, 2011; Bolnick *et al.*, 2011; Violle *et al.*, 2012). Genetic and phenotypic variation affects not only evolutionary but also ecological forces and factors including population dynamics, interaction strengths, and community composition (Allen *et al.*, 2018; Austin and Dunlap, 2019; Des Roches *et al.*, 2017; Frankham, 1996; Hausch *et al.*, 2018; Lloyd-Smith *et al.*, 2005; Start, 2019; Start and Gilbert, 2019; Vellend, 2006). These forces can change in time as the level of genetic variation in natural populations is not constant but can change on a much faster ecological time scale (Buckling and Rainey, 2002; Nijhawan *et al.*, 2019; Summers *et al.*, 2003). In particular, man-made events have lead to drastic changes in genetic variation (Jacquemyn *et al.*, 2009; Keller and Largiadèr, 2003; Mitrovski *et al.*, 2008; Smith *et al.*, 1991). Intraspecific variation is also expected to change as a result of coevolutionary processes. Character displacement, the divergence of mean phenotypes of two species in sympatry (Brown and Wilson, 1956; Dayan and Simberloff, 2005; Schulter and McPhail, 1992), is a well-studied example of temporal changes in phenotypes which decreases interspecific competition between the two species. Alternately, interspecific competition can be decreased by phenotypic diversification of one of the species (Abrams and Matsuda, 1994; Dieckmann and Doebeli, 1999; Winkelmann *et al.*, 2014). Overall, the importance of intraspecific variation in eco-evolutionary dynamics is well established. Thus, the description of the coevolutionary processes solely in terms of mean trait values may not be sufficient - one also needs to consider variances and higher-order moments of trait distribution.

Evolutionary theory has long been concerned with the problem of the maintenance of genetic variation (Barton, 1986; Clarke, 1979; Gavrilets, 2004; Gavrilets and Hastings, 1994b; Lande, 1975; Walsh and Lynch, 2018). There is now a rich variety of models explaining the maintenance of genetic variation by mutation-selection balance, frequency-dependent selection, spatial heterogeneity, *etc*. There is a number of theoretical tools for modeling coevolution of mean trait values. However significantly less efforts have focused on the dynamics of variances. In an early study of the dynamics of genetic variation, Bulmer (1971) used the infinitesimal model which assumes that a quantitative trait is controlled by infinitely many loci with infinitely small effects. In his model, selection builds linkage disequilibrium which changes genetic variation. Using a population genetics model with major loci, Gavrilets and Hastings (1994a, 1995) studied the dynamics of genetic variation under stabilizing selection. Finally, the adaptive dynamics approach has been used to study emergence of genetic polymorphism as a result of evolutionary branching (Dieckmann and Doebeli, 1999; Diekmann, 2004; Geritz *et al.*, 1998; Zu and Wang, 2013). All these methods have to deal with a trade-off between mathematical complexity and biological realism; each of these methods has its own advantages but also shortcomings. For example, Bulmer (1971) and Gavrilets and Hastings (1994a, 1995) ignored population densities, and the assumptions underlying the adaptive dynamics methods prevent one from exploring the changes in genetic variation in detail (Geritz *et al.*, 1998; Waxman and Gavrilets, 2005).

Here we develop an alternative framework for studying the effects of phenotypic variation in coevolution. Our framework is based on earlier single-species theoretical studies capturing both population densities and genetic variation. We apply our approach to three different types of two-species ecological interactions: (i) competition, (ii) exploiter-victim, and (iii) mutualism. For the cases of competition and exploiter-victim interactions, we study the conditions for coexistence, equilibrium trait distributions, and the relationship between the strength of interaction and phenotypic variance. In the case of mutualism, we study the conditions for equilibrium coexistence. Our framework relates stabilizing natural selection, trait-based interactions between coevolving species, phenotypic distributions, and population densities to understand the ecological consequences of intraspecific variation in two-species coevolutionary systems.

## Modeling framework

To introduce our approach, we start with the standard logistic model for the dynamics of the population size *N*(*t*) in time:

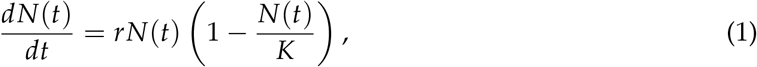

where *r* is the population growth rate at low densities and *K* is the carrying capacity. Here, the population size always approaches *K* asymptotically. This model implicitly assumes that all individuals are identical (Kot, 2001).

Roughgarden (1972) and Doebeli and Ispolatov (2010) extended this model for the case of heritable intraspecific trait variation. Following their method, we assume that individuals differ with respect to a single continuous trait *x*. Then the population density *ϕ*(*x*, *t*) of trait *x* at time *t* changes according to equation

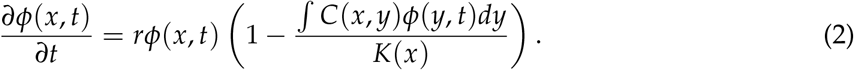

Here *r* is the growth rate at low densities (assumed to be independent of *x*), competition kernel *C*(*x*, *y*) measures the effect of competition with individuals with trait *y*, and function *K*(*x*) is the “carrying capacity” for individuals with trait value *x*. The total population size is given by the integral *N*(*t*) = *∫ϕ*(*y*, *t*)*dy*. Equation (2) implies that individuals reproduce asexually.

It is standard and mathematically convenient to use Gaussian functions 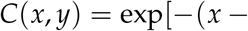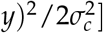 and 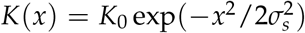. The former function implies that competition decreases with increasing the difference in trait values. The latter function assumes that carrying capacity decreases with deviation from the optimum trait value (which is set to zero without the loss of generality). Parameter *σ_c_* of the competition kernel measures a characteristic *range of competitive interference*: with small *σ_c_*, competition mostly happens between very similar organisms. Parameter *σ_s_* of the carrying capacity measures a characteristic *range of optimal trait values*: with small *σ_s_*, stabilizing selection is strong and only individuals with trait values close to the optimum can have high carrying capacity.

In this model, depending on parameter values the population evolves to one of two possible equilibrium states (Doebeli and Ispolatov, 2010). Specifically, if stabilizing selection is relatively strong, i.e. the range of optimal trait values is smaller than the range of competitive interference (*σ_s_ ≤ σ_c_*), the population becomes monomorphic with the optimum trait 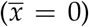 and the total equilibrium population density *N*^*^ = *K*_0_. This outcome is similar to that in the standard logistic model. However, if stabilizing selection is relatively weak (i.e., if *σ_s_ > σ_c_*), then the equilibrium distribution is Gaussian with the mean at the optimum 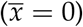 and a positive variance 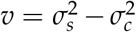 which increases with decreasing *σ_c_*. That is, if competition between dissimilar individuals is weak, variation can be maintained. The corresponding total equilibrium population density is 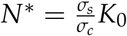 which is always larger than the total population density *K*_0_ at the monomorphic state. That is, maintaining genetic variation leads to larger population densities. Figure (1) illustrates these results.

**Figure 1:**
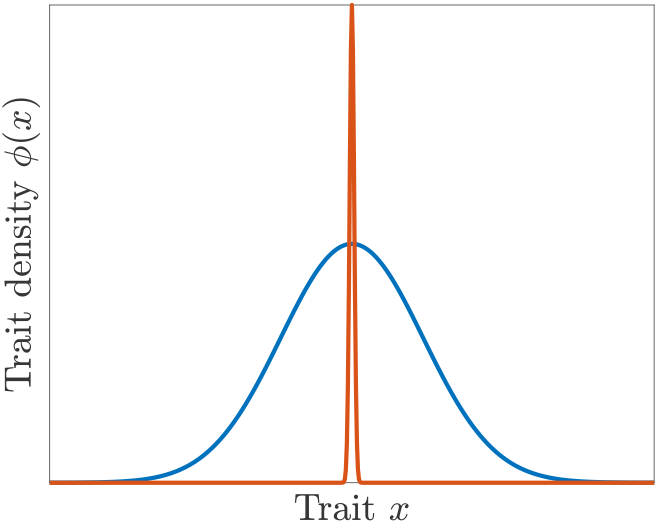
Equilibrium trait distributions in the single-species model. With stronger stabilizing selection 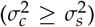, the population becomes monomorphic at the optimum trait value *x* = 0 (orange curve). With stronger competition 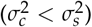, the equilibrium trait distribution is Gaussian with a positive variance 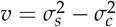 (blue curve).

Below we will generalize this approach for three different two-species models which we will study using analytical approximations and numerical solutions (see the Supplementary Information for details on our numerical method).

## Two-species competition

The standard Lotka-Volterra competition model describes the dynamics of two competing species with densities *N*_1_ and *N*_2_:

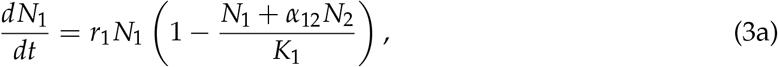

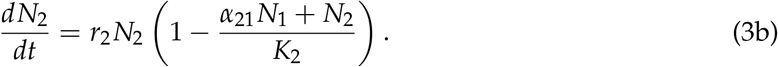

Here, for species *i* (*i* = 1, 2), *r_i_* is the growth rate at low densities, *K_i_* is the carrying capacity in the absence of the competing species, and parameters *α*_12_ and *α*_21_ represent the strength of between-species competition relative to that within species. In this model, the necessary and sufficient condition for coexistence is that within-species competition is stronger than between species competition for both species: *α*_12_*K*_2_/*K*_1_ < 1 and *α*_21_*K*_1_/*K*_2_ < 1. If the condition is satisfied in one species (say *α*_12_*K*_2_/*K*_1_ < 1) and not satisfied in the other species (so that *α*_21_*K*_1_/*K*_2_ ≥ 1) then the first species survives and the second species goes extinct. We then say that the first species *1* is a stronger competitor and species 2 is a weaker competitor. If *α*_12_*K*_2_/*K*_1_ ≥ 1 and *α*_21_*K*_1_/*K*_2_ ≥ 1, then one species survives and the other becomes extinct based on initial conditions.

We extend the Lotka-Volterra competition model to individuals differing in continuous traits *x* in the first species and *y* in the second species by adapting the single species approach described above:

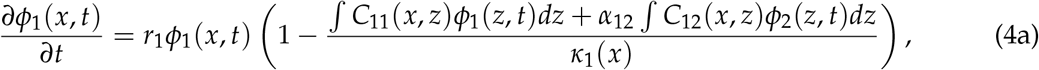

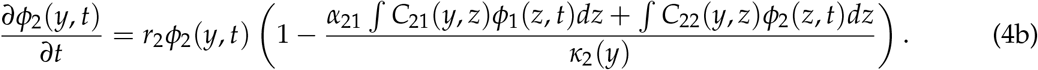

Here *ϕ_i_*, *r_i_* and *κ_i_* are the population density of the trait, the intrinsic growth rate, and carrying capacity for species *i*, and *C_ij_* are the corresponding competition kernels. As above, we assume that carrying capacity, intraspecific competition, and interspecific competition kernel functions are Gaussian: 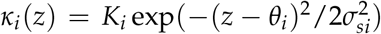, 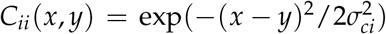, and 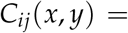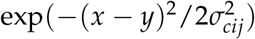 respectively. Here *θ_i_*, *σ_ci_*, and *σ_si_* are the optimum trait value, the range of within-species competitive interference, and the range of optimal trait values for species *i*, *σ_c_*_12_ and *σ_c_*_21_ measure the ranges of between-species competitive interference, whereas *α*_12_ and *α*_21_ measure the strength of interspecific competition due to trait-independent between-species differences.

### Results

To find sufficient conditions for coexistence we used mutual invasibility analysis (Armstrong and McGehee, 1980; Geritz *et al.*, 1998). The idea underlying this method is that the two species will coexist only if each of them can invade a resident population of the other species at equilibrium. For example, in the Lotka-Volterra competition model described above, species 2 can invade a resident population of species 1 at equilibrium (*N*_1_ = *K*_1_) from low population density (*N*_2_ ≈ 0) only when 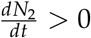. This gives the invasion criterion for species 2: *α*_21_*K*_1_/*K*_2_ < 1. Invasion criteria are sufficient conditions for coexistence since they guarantee neither species can go extinct.

Assume first that the resident population is monomorphic which is the case if stabilizing selection is sufficiently strong (*σ_sj_* ≤ *σ_cj_*). Then the invasion criterion for the invader species *i* is identical to that in the standard Lotka-Volterra competition model:

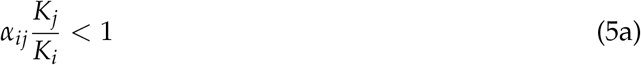

(see Appendix A for details). Assume next that the resident population is polymorphic which is the case if *σ_sj_ ≥ σ_cj_*. From the single-species model, the variance of a polymorphic resident population is 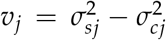 and population size is 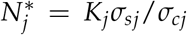. In this case, if stabilizing selection in the invader is weak enough 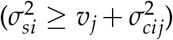, it will invade always. Otherwise, invasion happens whenever

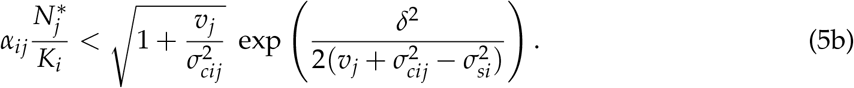

where *δ* = |*θ_i_* − *θ_j_*| is the difference between the optimum trait values. Note that increasing the difference in the optimum trait values *δ*, decreasing the strength of stabilizing selection in the invader (i.e., increasing *σ_si_*), or decreasing the range of between-species competitive interference *σ_cij_* always increases the right-hand side of equation (5b) and, thus, makes invasion easier. If *δ* = 0, increasing the ratio 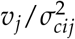, i.e. decreasing the range of between-species competitive interference relative to the phenotypic variance in the resident species, makes invasion easier. Figure (2) illustrates our analytical results.

**Figure 2:**
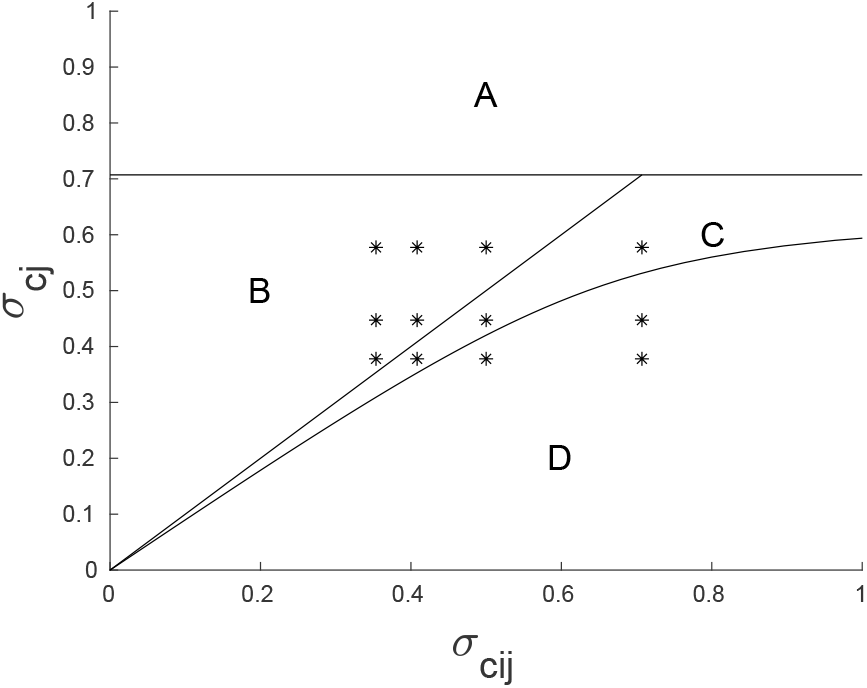
Conditions for invasion as a function of the ranges of between-species *σ_cij_* and within-species *σ_cj_* competitive interference. In region A, *σ_cj_ > σ_sj_*, the resident is monomorphic, and the invasion condition is based on inequality (5a). The line separating regions B and C is based on whether stabilizing selection on the invader is sufficiently weak to guarantee invasion (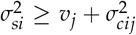 in region B). In region B, invasion is guaranteed for any choice of *α_ij_*. Inequality (5b) determines the boundary between region C and D. It is satisfied in region C and not satisfied in region D. The black dots corresponds to the parameter choice for the numerical simulations illustrated in Figure (3) below. Other parameters: *σ_sj_* = 0.71, *δ* = 0, *α_ij_* = 0.9.

The invasion analysis provides only sufficient conditions for coexistence and does not tell anything about the equilibrium distributions of traits. Therefore we supplement it with a numerical study of the model dynamics for different combinations of parameters and initial conditions using an adaptive finite difference method (see Appendix B, also Doebeli (2011)). We describe our results next. In all cases considered, the system evolves to an equilibrium. If only one species survives, the equilibrium trait distribution matches the one predicted by equation (2).

At a coexistence equilibrium with both species polymorphic there are three possibilities: (i) both species are unimodal, (ii) one species is unimodal and another bimodal, or (iii) both species are bimodal. These outcomes are illustrated in Figure (3). The values of parameters *σ_cij_* and *σ_cj_* used in these graphs are marked by dots in Figure (2). Assuming that species 2 can invade a resident population of species 1 (i.e. that inequality (5b) holds for *i* = 2, *j* = 1), parameters relevant for species 1’s invasion may lie in one of the regions A-C. In region B, species 1 is bimodal if the range of within-species competitive interference in species 2 is large and unimodal otherwise. Species 2 is bimodal only if the range of within-species competitive interference in species 2 is intermediate. In region A and C, both species are unimodal.

**Figure 3:**
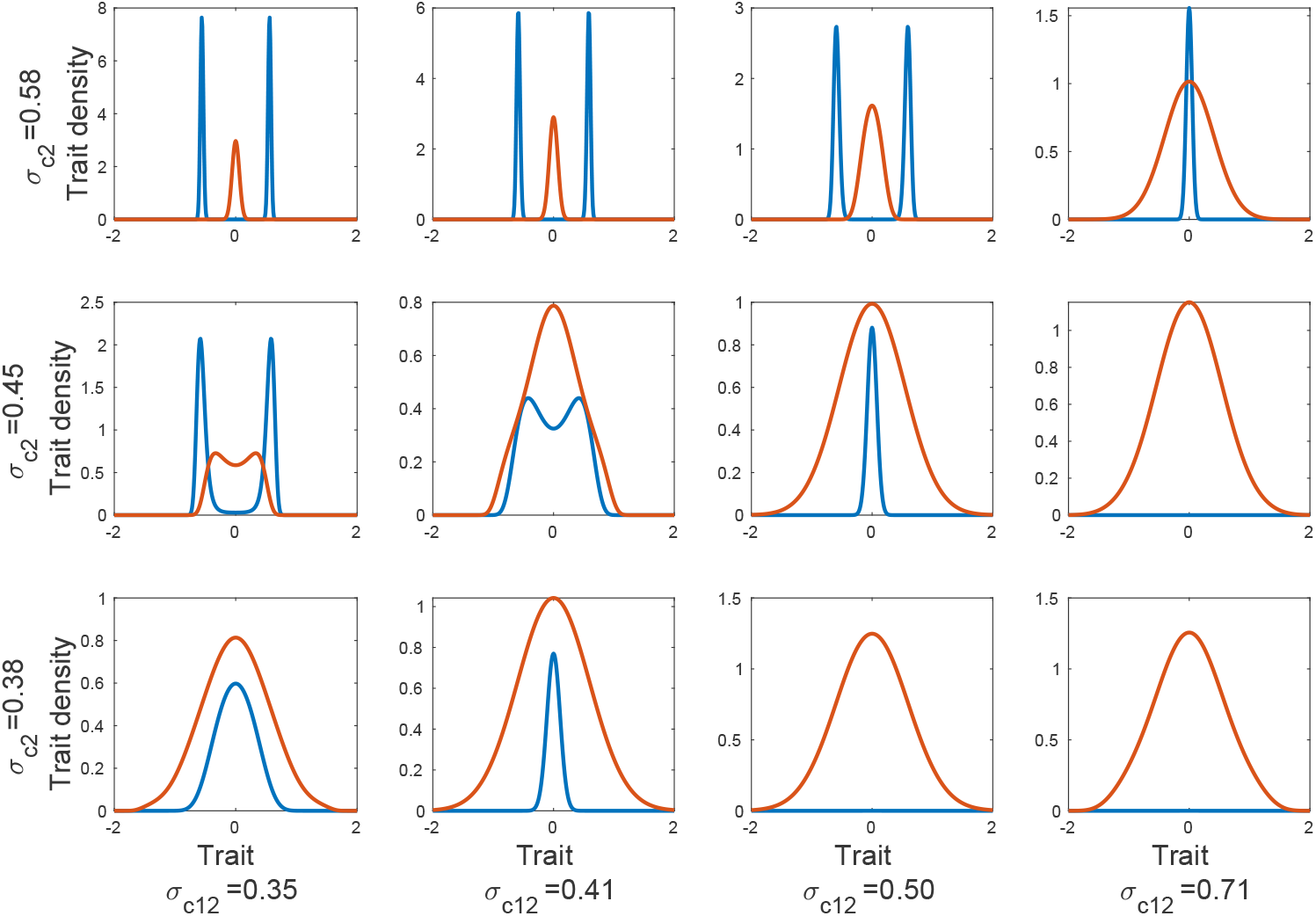
Equilibrium trait distributions of species 1 (blue) and species 2 (orange) from numerical simulations for parameters values *σ_cij_* and *σ_cj_* marked by dots in Figure (2). Identical initial phenotypic distributions 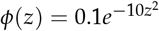. For all the simulations, *α*_12_ = 0.9, *α*_21_ = 0.8, *σ_c_*_1_ = 0.71, *σ_c_*_21_ = 0.71, *σ_s_*_1_ = 0.71, *σ_s_*_2_ = 0.71, *δ* = 0, *r*_1_ = 1, *r*_2_ = 1, *K*_1_ = 1, *K*_2_ = 1.

In the single species model, the equilibrium population size and variance decrease with the range of within-species competitive interaction *σ_ci_* (see above). To explore the effects of *σ_ci_* in the case of two coexisting species, we varied *σ_c_*_2_ assuming that parameters for species 1 are in regions B or C, and for species 2 in region A of Figure (2). Figure (4) shows the equilibrium variance and population when the both species coexist (solid lines) and the analytical solution of the single-species model (dotted line). We find that the equilibrium variance of species 2 decreases with *σ_c_*_2_ and is always smaller when the other species is present. Equilibrium variances of the two species are inversely related. Similarly, the equilibrium population size of species 2 decreases with *σ_c_*_2_ whereas that of species 1 increases with *σ_c_*_2_. Equilibrium population sizes of both the species are lower than their respective single-species equilibrium.

**Figure 4:**
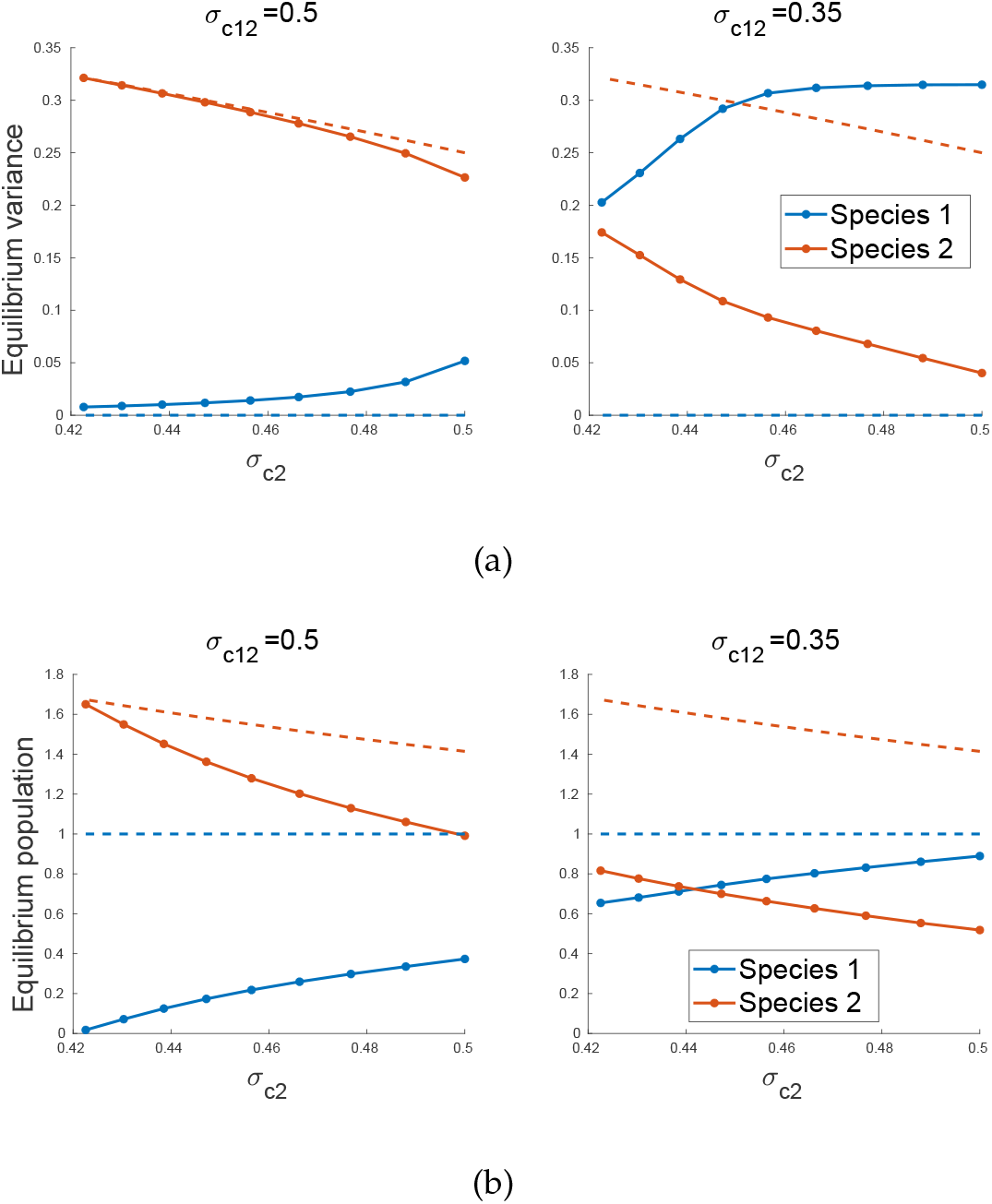
Effect of the range of within-species competitive interference *σ_c_*_2_ on the equilibrium variance and population sizes of the two species. The dotted lines are based on the equilibrium of the single-species model and the solid lines are based on numerical solutions of the two-species model with identical initial phenotypic distributions *ϕ*(*z*) = 0.1 exp(−10*z*^2^). The left and right panels correspond to coexistence in region C and B in Figure (2), respectively. Other parameters: *α*_12_ = 0.9, *α*_21_ = 0.8, *σ_c_*_1_ = 0.71, *σ_c_*_21_ = 0.71, *σ_s_*_1_ = 0.71, *σ_s_*_2_ = 0.71, *δ* = 0, *r*_1_ = 1, *r*_2_ = 1, *K*_1_ = 1, *K*_2_ = 1.

## Exploiter-victim interactions

Here we consider exploiter-victim interactions such as between a predator and a prey or a parasite and a host (Clayton *et al.*, 1999; Davies and Brooke, 1988; Gavrilets, 1997; Soler *et al.*, 2001; Toju and Sota, 2006). Writing the population density of the victim species as *N*_1_(*t*) and that of the exploiter species as *N*_2_(*t*), and assuming that the exploiter has an obligate relationship with the victim, we start with the model

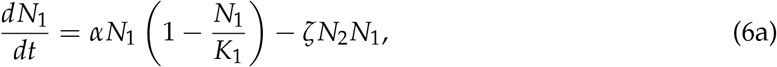

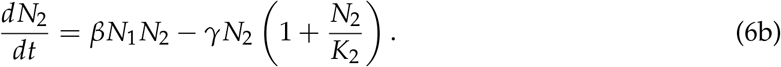

Here, the intrinsic growth rate of the victim is *α*, the intrinsic death rate of the exploiter is *γ*, *β* and *ζ* are the exploiter birth rate and victim death rate due to exploitation, *K*_1_ and *K*_2_ are characteristic population densities. This is a generalization of the classical Lotka (1920) model to which we have added a quadratic death rate term due to within-species competition. [This change also allows one to avoid structural instability inherent in the Lotka-Volterra model (Kot, 2001). We recover the Lotka-Volterra model in the limit of large *K*_1_ and *K*_2_.]

In this model, both species coexist at an asymptotically stable equilibrium if

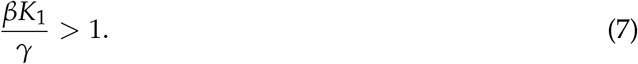

The numerator of the above ratio is the growth rate of the exploiter when the victim is at carrying capacity (i.e., *N*_1_ = *K*_1_) and the denominator is the exploiter’s death rate at low densities. The corresponding equilibrium densities are 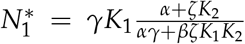 and 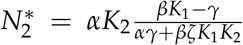. If the inequality above is reversed, only the victim species survives. This model does not lead to exploiter-victim cycles.

We extend the above model to individuals differing in continuous traits *x* in the victim and *y* in exploiter:

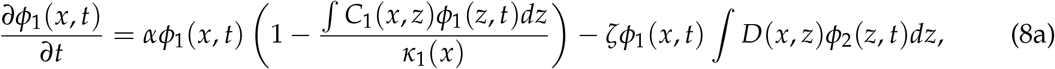

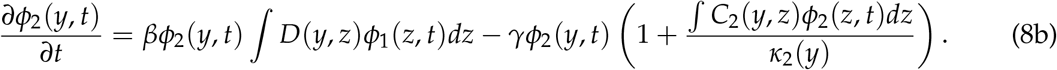

Here *ϕ*_1_(*x*, *t*) and *ϕ*_2_(*y*, *t*) are the corresponding densities of the traits, and parameters *α*, *β*, *γ* and *ζ* have the same meaning as above. Similar to the competition model with continuous traits, we assume that selection and within-species competition kernels are Gaussian: 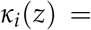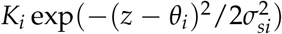 and 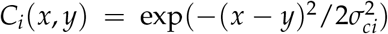, where parameters *θ_i_*, 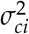 and 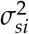 have the same meaning as before. Assuming that exploiter-victim interactions are based on trait matching (Gavrilets, 1997; Yoder and Nuismer, 2010), the exploitation kernel can be modelled as a Gaussian function: 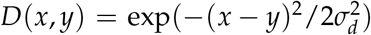, where 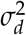 measures the *range of exploitative interactions*. For example, with small 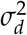 the exploiter can utilize only victims with very similar trait values.

### Results

Using the invasibility analysis, for the exploiter to coexist with the victim, it should be able to grow from small densities. Consider first the case where stabilizing selection in the victim is strong (*σ_c_*_1_ ≥ *σ_s_*_1_). In this case, the victim is monomorphic and the sufficient conditions for coexistence is identical to the coexistence condition from the model with no individual variation (inequality (7)). In contrast, if *σ_c_*_1_ *< σ_s_*_1_, the victim is polymorphic with equilibrium variance 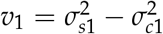 and equilibrium population size 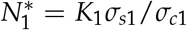 in the victim-only model. In this case, the sufficient condition for a successful invasion of the exploiter (and thus for coexistence) is

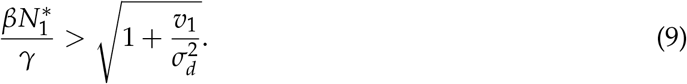

In particular, this shows that increasing the range of exploitative interactions *σ_d_* simplifies survival of the exploiter.

We did not observe any non-equilibrium dynamics in numerical simulations. The equilibrium trait distribution of the victim matched the single-species equilibrium when the exploiter did not survive. Figure (5) illustrates trait distributions when the species coexist. The distributions are unimodal if the exploiter death rate (*γ*) is large. If it is small, the victim diversifies around the exploiter to survive. [This situation was dubbed the Burridan’s Ass regime in Gavrilets and Waxman (2002).] If stabilizing selection in the victim (*σ_c_*_1_) is weak, the diversification in the victim can be followed by that in the exploiter (top right graph in Figure (5)).

**Figure 5:**
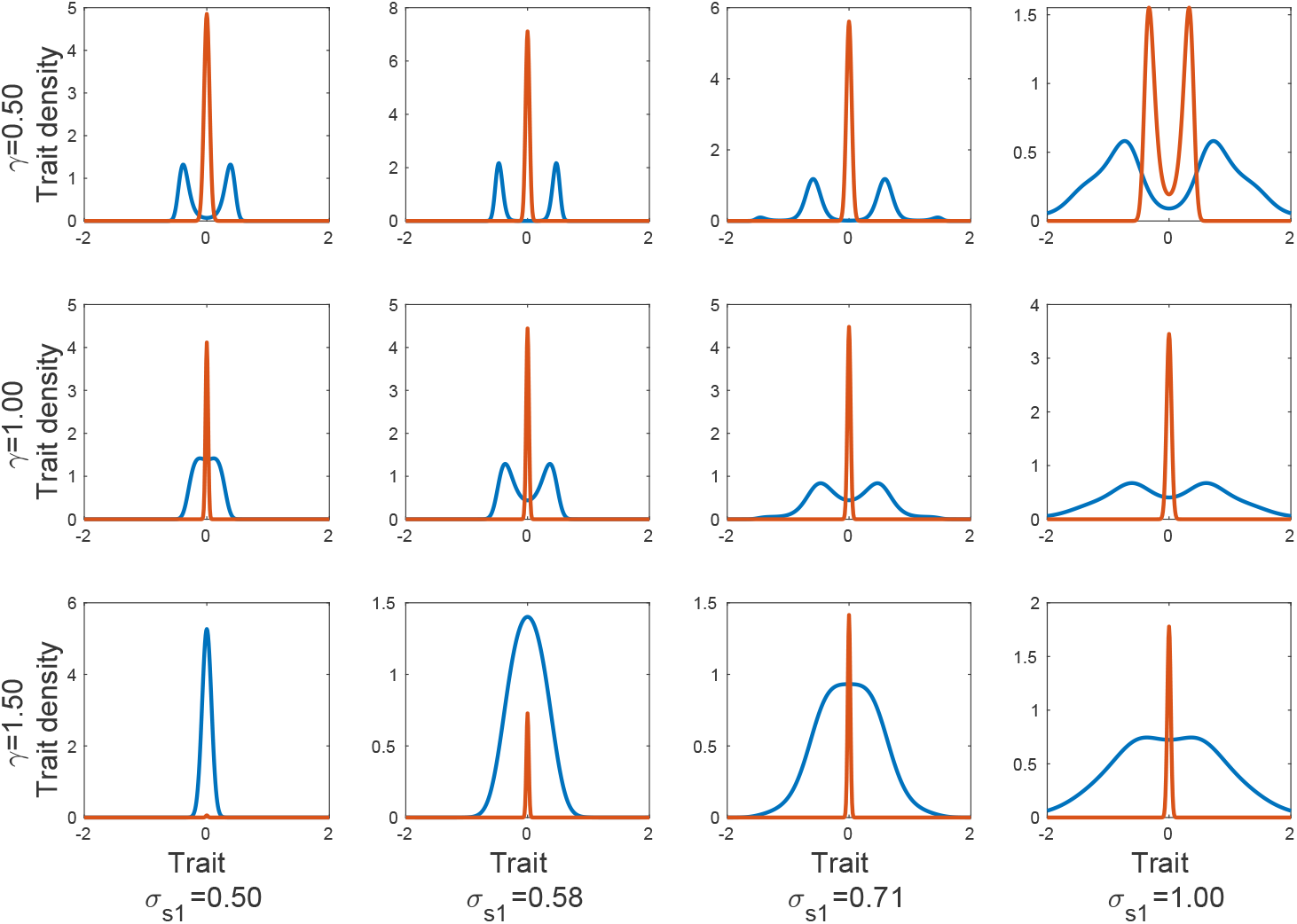
Examples of equilibrium trait distributions of the exploiter (orange) and victim (blue) from numerical simulations with identical initial trait distributions *ϕ*(*z*) = 0.1 exp(−10*z*^2^). Other parameters: *α* = 1, *β* = 1.5, *ζ* = 1, *σ_c_*_1_ = 0.5, *σ_c_*_2_ = 0.71, *σ_d_* = 0.71, *δ* = 0, *K*_1_ = 1, *K*_2_ = 1.

The equilibrium variance in both species increases with the strength of competition among victims (Figure (6)). The exploiter’s variance is smaller than that in the victim even when the exploiter trait distribution is bimodal (*σ_s_*_1_ = 1, *γ* = 0.5 in Figure (5)). Figure (6) also shows that if the exploiter survives, the victim’s variance is larger and the population size is smaller than the equilibrium values of the victim-only model.

**Figure 6:**
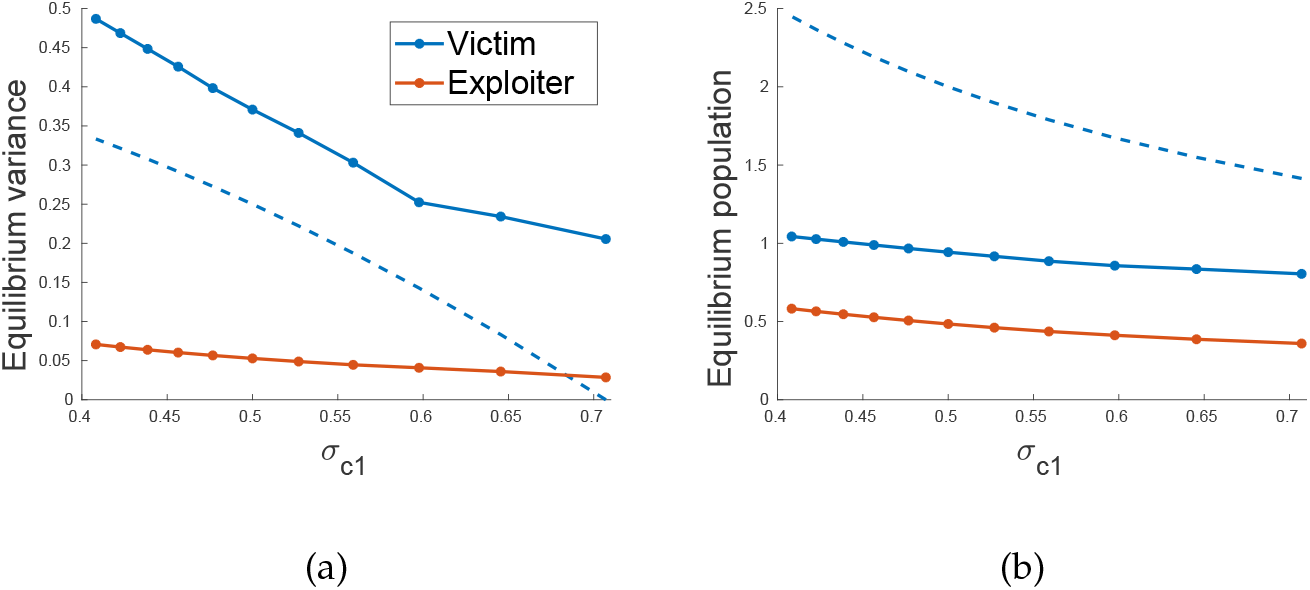
The equilibrium variances and population sizes of both species when the exploiter survives. The dotted line is the equilibrium variance of the victim in the single-species model and the solid lines are numerical simulations using identical phenotypic distribution for the victim and exploiter (*ϕ*(*z*) = 0.1 exp(−10*z*^2^)). Other parameters: *α* = 1, *β* = 2, *ζ* = 1, *γ* = 1, *σ_c_*_2_ = 0.32, *σ_s_*_1_ = 0.71, *σ_s_*_2_ = 0.71, *δ* = 0, *K*_1_ = 1, *K*_2_ = 1.

## Mutualism

In this section we consider mutualistic interactions where both species benefit from the interaction (Bronstein, 2015). We start with a classical mutualism model (Gause and Witt, 1935; Kostitzin, 1939) describing the dynamics of a pair of mutualistic partners with densities *N*_1_ and *N*_2_:

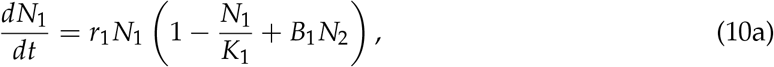

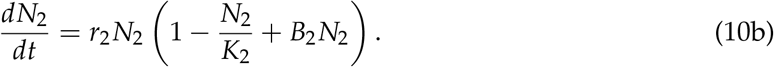

Here, the mutualistic benefit for species *i* is denoted by *B_i_* and the carrying capacity is denoted by *K_i_*. A stable equilibrium exists if and only if *B*_1_*B*_2_*K*_1_*K*_2_ < 1. At this equilibrium, the population size of species *i* is 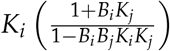. If the above inequality does not hold, both species grow to infinite sizes. This unbounded growth is sometimes referred to as the “orgy of beneficial mutualism” (May, 1981).

Allowing for individual differences in continuous traits *x* in the first species and *y* in the second species, the corresponding dynamics of the mutualistic system are described by equations

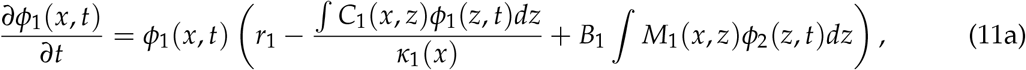

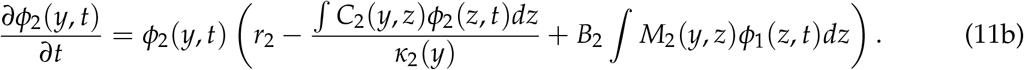

Here *ϕ_i_*, *r_i_*, and *κ_i_* represent the population density of the trait, the intrinsic growth rate, carrying capacity for species *i*, and *C_i_* are the corresponding competition kernels. Similar to the other models, we assume carrying capacity and competition kernel functions are Gaussian: 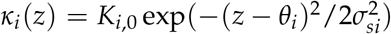 and 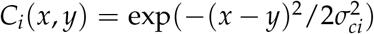. Assuming that mutualism is based on trait matching (Brouat *et al.*, 2001; Yoder and Nuismer, 2010), the mutualism kernel *M_i_* can be modelled by a Gaussian function: 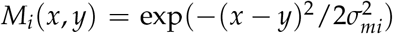, where *σ_mi_* measures the *range of mutualistic interactions*, *i.e.* range of phenotypes over which an individual receives mutualistic benefits. Parameters *B_i_* represent mutualistic benefits due to other trait independent factors.

### Results

Similar to the case of no intraspecific variation, in this model the two species either reach an equilibrium with finite population sizes or grow to infinite populations due a positive feedback of mutualistic benefits. To find the conditions for the population to reach an equilibrium, an approach similar to the invasion analysis does not work. Instead we first find upper bounds for the time series of population sizes of the two species, and then determine conditions for the upper bound to converge. This gives us the necessary conditions for stable coexistence (see Appendix A for details).

Our results are as follows. If stabilizing selection is sufficiently strong (*σ_ci_ ≥ σ_si_*) in both species, then both species are monomorphic and the necessary condition for an equilibrium is identical to that in the model with no intraspecific variation:

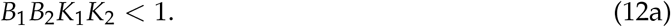

If stabilizing selection is strong in only one of the species (say, 2nd), then the necessary condition for an equilibrium is

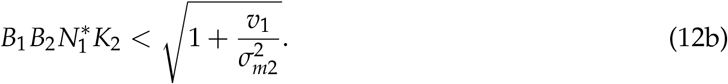

Here 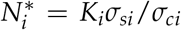 and 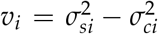 are the equilibrium population size and variance of the single-species model for species 1. This shows that if stabilizing selection is strong in one of the species, an equilibrium is easier to achieve when the range of mutualistic interactions (*σ_m_*_2_) is small.

Finally, if stabilizing selection is weak in both species, then the necessary condition for an equilibrium is

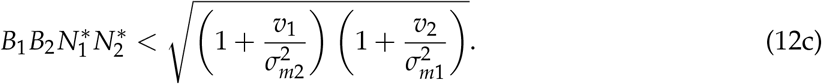

In contrast to the case of strong stabilizing selection in only one of the species, here an equilibrium is easier to achieve when the range of mutualistic benefits is small in either of the two species. Overall, if variation is maintained (i one of both species), it is less likely that their population growth would be unbounded.

Numerical exploration of the model revealed two types of polymorphic equilibrium trait distributions: (i) a unimodal (Figure (7(a))), or (ii) a multimodal (Figure (7(b))). In the multimodal case, there is one large peak and two smaller peaks (Figure (7(b))) or one large and one small peak. The latter case is observed when *δ* is large and the former when *δ* is small. These additional smaller peaks exist because with large range of within-species competition *σ_ci_*, mutualistic benefits do not decrease as quickly as competitive costs (decrease in growth rate due to competition) for traits farther away from the mean of the distribution.

**Figure 7:**
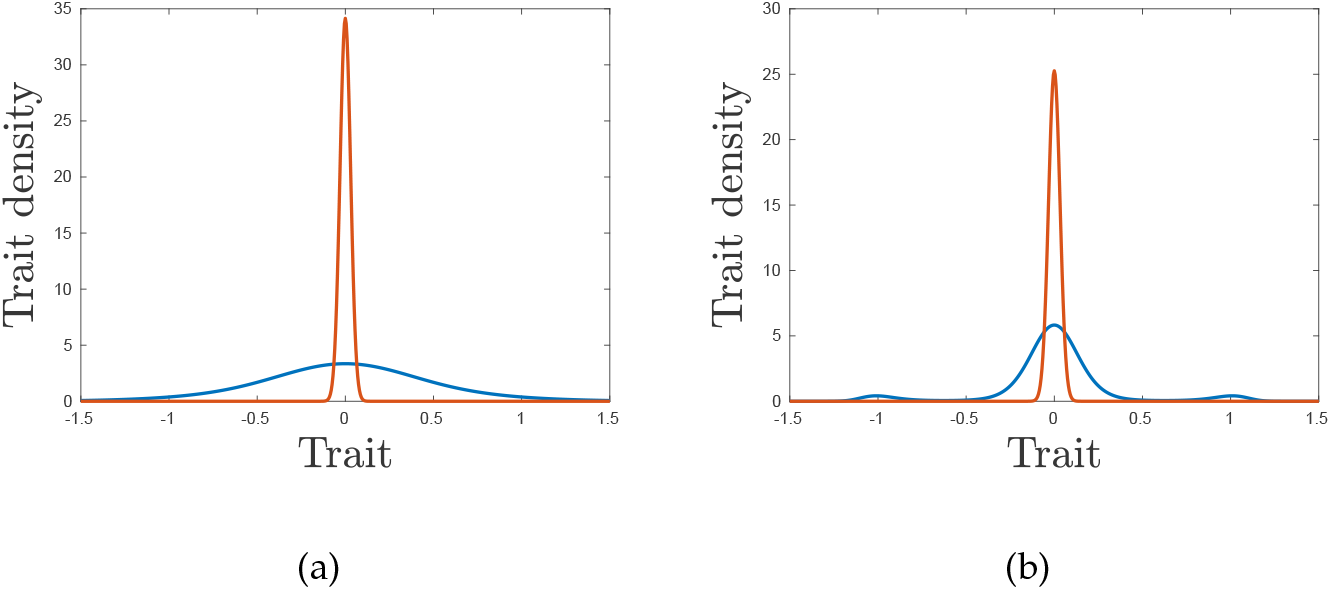
Common equilibrium trait distributions of mutualists. (a) *σ_c_*_1_ = 0.35 and (b) *σ_c_*_1_ = 0.53. Species 1 (blue) and mutualist 2 (orange); identical initial trait distributions, other parameters *ϕ*(*z*) = 0.1 exp(−10*z*^2^) with *r*_1_ = 1, *r*_2_ = 1, *B*_1_ = 0.5, *B*_2_ = 0.5, *σ_c_*_2_ = 0.71, *σ_s_*_1_ = 0.71, *σ_s_*_2_ = 0.71, *σ_m_*_1_ = 0.71, *σ_m_*_2_ = 0.71, *δ* = 0, *K*_1_ = 1, *K*_2_ = 1. For *δ* > 0, the mean of the distributions will shift but the shape remains the same.

We also found that as the range of within-species competition decreases, the equilibrium variance increases (Figure 7(a)). Figure (8) show that the equilibrium variance increases with smaller range of within-species competition irrespective of the range of within-species competition in the other species. Also, the equilibrium variance is always smaller than the equilibrium variance of the single-species model. In comparison to the single-species model, the equilibrium populations are higher when the mutualist partner is present. The equilibrium population becomes larger as the range of within-species competition in either species decreases.

**Figure 8:**
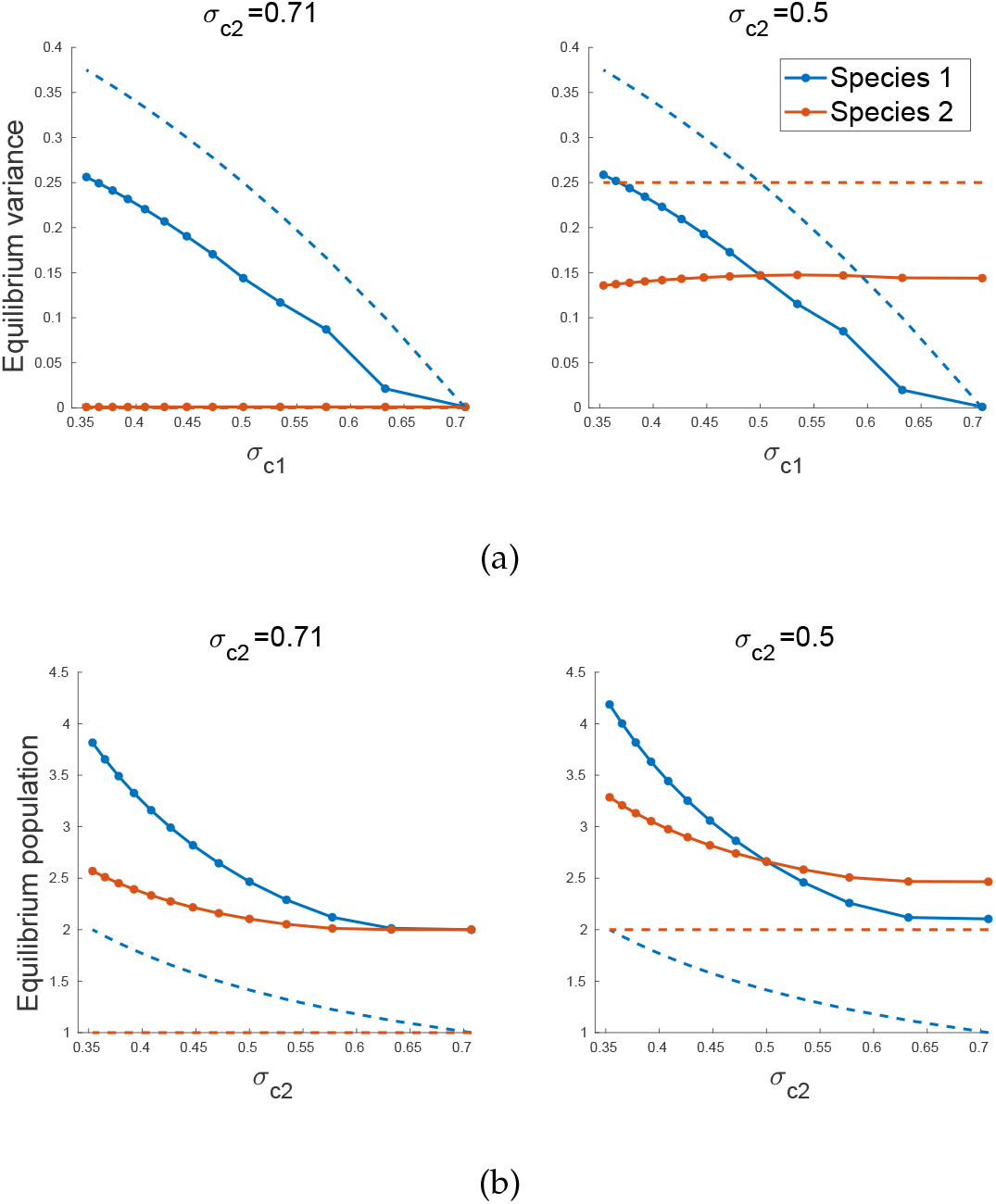
Effect of the range of within-species competitive interference on equilibrium variance and population of the mutualists. The dotted lines show the equilibrium of a single-species model and the solid lines are based on the numerical solution of the two species model with initial phenotypic distributions 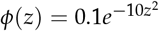. Other parameters: *r*_1_ = 1, *r*_2_ = 1, *σ_s_*_1_ = 0.71, *σ_s_*_2_ = 0.71, *σ_m_*_1_ = 0.71, *σ_m_*_2_ = 0.71, *B*_1_ = 0.5, *B*_2_ = 0.5, *δ* = 0.

## Discussion

Although the dynamic patterns of between-species interactions are expected to strongly depend on intraspecific variation, how exactly ecological and evolutionary processes interact is still largely an open question. Here we approached this question theoretically using three simple two-species models describing competition, exploiter-victim interaction, and mutualism. In our models, individuals differ with respect to a single quantitative character which controls both within- and between-species density-dependent interactions and, simultaneously, is subject to stabilizing natural selection. We analysed conditions for species coexistence, equilibrium population densities as well as the characteristics of trait distributions observed at equilibrium.

For intraspecific variation to be important, it needs to be maintained. Our results show that without mutation, intraspecific variation is lost if stabilizing selection is strong enough, specifically, if the range of optimal traits values is narrower than the range of within-species competitive interference (*σ_s_ < σ_c_*). In this case, the outcomes of population dynamics are the same as predicted by standard ecological models neglecting intraspecific variation. The condition *σ_s_ < σ_c_* for the loss of intraspecific variation is the same as the one in single-species models (Doebeli and Ispolatov, 2010; Roughgarden, 1972). Intraspecific variation can be maintained if stabilizing selection is weak enough in at least one species of the pair. In the discussion below, we assume that this is the case.

Consider first between-species competition. In classical ecological models of competition, there are three possible outcomes: the extinction of a weaker competitor and persistence of a stronger competitor, survival of one species or another depending on initial densities, or coexistence. With intraspecific variation maintained, these three possible outcomes are still possible but the conditions for them to be observed depend on parameters characterizing ranges of interference. In particular, small ranges of between-species interference (*σ_cij_*) make coexistence a more likely outcome. When the species coexist, their equilibrium trait distribution can be unimodal for both species, bimodal for both species, or unimodal for one species and bimodal for the other. The weaker competitor (based on ecological model) has a bimodal distribution when the range of between-species competition is large, and unimodal otherwise. The strong competitor has a unimodal trait distribution when the range of between-species competition is small or large, and a bimodal distribution for a narrow range of intermediate values.

Second, in classical ecological models of exploiter-victim interactions there are two possible outcomes: the victim species survives and the exploiter is extinct, or coexistence. These outcomes are also possible if within-species variation is maintained. In general, large ranges of exploitative interactions (*σ_d_*) promotes survival of the exploiter. When the species coexist, their equilibrium trait distributions can be both unimodal, both bimodal, or unimodal in one species and bimodal in the other. When the exploiter’s death rate (*γ*) is high, both the exploiter and victim have unimodal trait distributions at the coexistence equilibrium. When the exploiter’s death rate is low, the victim diversifies and its equilibrium trait distribution is bimodal. If stabilizing selection in the victim is weak, its trait distribution becomes bimodal which can be followed by the evolution of bimodality in the exploiter. Third, in classical ecological models of mutualism there are two possible outcomes: the two species coexist at finite population sizes, or both species grow indefinitely due to non-diminishing mutualistic benefits. With intraspecific variation maintained, small range of mutualistic interactions (*σ_m_*) promotes coexistence at finite sizes. The equilibrium trait distribution is typically unimodal, but can become multimodal (with two or three peaks) if stabilizing selection is strong enough. In this case, only one of the peaks has a high trait density, while other peak(s) have a much smaller trait density.

In single-species models allowing for heritable intraspecific variation (such as given by equation (2)), the appearance of bimodal trait distributions or evolutionary branching require a non-Gaussian competition kernel, e.g. an asymmetric (Kisdi, 1999) or a platykurtic (Doebeli and Ispolatov, 2010). In contrast, our coevolutionary models show that species interactions could lead to bimodal trait distribution even for Gaussian competition and interaction functions. In our competition model, the equilibrium trait variances are smaller than those in the corresponding single-species model. For the exploiter-victim interaction, the equilibrium trait variance of the exploiter is always much smaller than that in the victim. In comparison to the single-species model, the victim always has a higher variance in presence of the exploiter. In the case of mutualism, equilibrium trait variances are smaller for both mutualist species compared to those in the corresponding single-species model. Overall, we find that coevolutionary interactions lead to smaller trait variances except for the victims in exploiter-victim interactions.

Both theoretical and empirical studies have explored the ecological consequences of intraspecific variation (Austin and Dunlap, 2019; Breza *et al.*, 2012; Des Roches *et al.*, 2017; Frankham, 1996; Gavrilets, 1997; Hart *et al.*, 2016; Lichstein *et al.*, 2007; Start, 2019; Start and Gilbert, 2019). However only few earlier theoretical studies allowed for within-species variation to evolve (Kopp and Gavrilets, 2006; Nuismer *et al.*, 2005). An interesting consequence of heritable trait variation is character displacement when between-species competition is reduced due to the divergence of mean phenotypes (Brown and Wilson, 1956; Dayan and Simberloff, 2005; Schulter and McPhail, 1992). Our results suggest competition can also be reduced due to an increase in phenotypic variances, or when the distributions become multimodal.

Our models show that the relationship between the range of optimum traits (which depends on the strength of stabilizing selection) and the ranges of within- and between-species interactions is an important determinant of coevolutionary dynamics. In general, a larger range of optimum traits relative to the range of within-species interactions leads to the maintenance of trait variation which in turn allows for competitors to coexist, exploiters to survive, and mutualists to reach a stable equilibrium. Increased evolutionary flexibility allowed for by intraspecific variation potentially offers a way to reconcile differences in empirical observations, some of which shows that intraspecific variation promotes coexistence (Clark, 2010; Fricke *et al.*, 2019; Jung *et al.*, 2010), while others suggest that it restricts it (Hausch *et al.*, 2018). The coexistence conditions we have derived extend coexistence theory (Barabás *et al.*, 2018; Chesson, 2000; Ellner *et al.*, 2019) by including the effect of heritable intraspecific variation, stabilizing selection, and trait-dependent competition. Our models also demonstrate that with heritable intraspecific variation maintained, the strength of trait-based interactions can change through time.

Cyclical dynamics in exploiter-victim interactions have been of great interest in ecological (Lotka, 1920; Turchin, 2003) and evolutionary (Gavrilets, 1997; Kopp and Gavrilets, 2006; Nuismer and Doebeli, 2004; Nuismer *et al.*, 2005) models. However cycling was not possible in our basic model (equations (6)) and we did not observe it in our extension of that model for the case of within-species variation. We note that Nuismer *et al.* (2005) investigated a range of evolutionary models of victim-exploiter type and concluded that cycling happens only under certain conditions.

Our results show that exploiter-victim interactions and competition can lead to bimodal trait distributions. Such distributions can emerge via the process of evolutionary branching (Geritz *et al.*, 1998) and can potentially lead to speciation. Doebeli and Dieckmann (2000) studied coevolutionary interaction between an exploiter and a victim using adaptive dynamics, and found that a large range of exploitative interaction and weak stabilizing selection in the victim can lead to evolutionary branching in exploiters. We find similar relationships between the range of exploitative interaction, the strength of stabilizing selection in the victim, and bimodality of the exploiter’s trait distribution when the exploiter’s death rate is high. Yoder and Nuismer (2010) studied phenotypic diversification in a metapopulation. They found that in coevolutionary interactions where fitness of at least one species is reduced when its traits match the other species, phenotypic diversity is higher compared to diversification by spatially variable selection without coevolution. We find that such costly trait matching leads to a higher trait variance in the victim, but can lead to a lower variance for a competitor. This is due to one of the competitor diversifying around the other competitor (*i.e.* bimodal distribution with the modes on either side of the mean trait of the other species), and competing from both directions of the mean trait value of the other species (see Figure (3), *σ_c_*_2_ = 0.58, *σ_c_*_12_ = 0.35).

The main limitations to our approach in terms of biological realism are that we ignored mutation and sexual reproduction. Including mutation would make it easier to maintain intraspecific variation at low levels but is not expected to significantly change our results. For analytical convenience, we assumed clonal reproduction in all our models neglecting the homogenizing force of sexual reproduction. Earlier theoretical studies of coevolution find that genetic details can lead to novel dynamics (Kopp and Gavrilets, 2006; Nuismer and Doebeli, 2004; Nuismer *et al.*, 2005). Some of the effects we observed (e.g., bimodal trait distributions) may be an artifact of our simplifying assumptions. Future work should focus on extending these models to sexually reproducing populations. The analytical conditions we have found are only sufficient (but not necessary) in the case of competition and exploiter-victim interaction, and necessary (but not sufficient) in the case of mutualism.

Our work adds to the toolkit of theoretical studies of coevolution a numerical approach (described in the SI) for modeling the dynamics of population densities and phenotypic distributions under different two-species interactions. We have also developed a novel application of the invasion analysis (Armstrong and McGehee, 1980), and derived an approximate analytical condition for existence of a stable coexistence equilibrium for mutualists (Appendix A). Our methods allowed us to obtain analytical results without the common assumption of weak selection in co-evolutionary models (Gavrilets, 1997; Nuismer and Doebeli, 2004). Overall, our approach allows us to understand better the role of heritable trait variation in coevolutionary systems, which is the first step towards achieving a better understanding of the implications of heritable trait variation in evolutionary community ecology (McPeek, 2017).

## Acknowledgements

We thank E. Derryberry, J. Day, D. Tverskoi, B. Fitzpatrick, and J. Bailey for comments and suggestions. Supported by the Office of Naval Research grant W911NF-17-1-0150, the National Institute for Mathematical and Biological Synthesis through NSF Award #EF-0830858, and by the University of Tennessee, Knoxville.

## Appendix A: Analytical solution

## Competition

Based on mutual invasibility analysis, we can derive conditions for each species to grow from low density in a population of the other species. In the absence of the species 1, species 2 will reach the equilibrium of the one species model (equation (2)). If 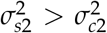, the trait distribution at the single-species equilibrium will be 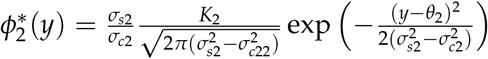. Else, it would be a delta function at *θ*_2_. Invasion criteria for species 1 in that case turns out to be identical to the population dynamics model (*i.e., α*_12_ *< K*_1_/*K*_2_).

For species i to coexist with species j, atleast individuals of some trait *x* needs to survive when *ϕ*_1_(*x*) ≈ 0 and 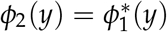. From equation (4),

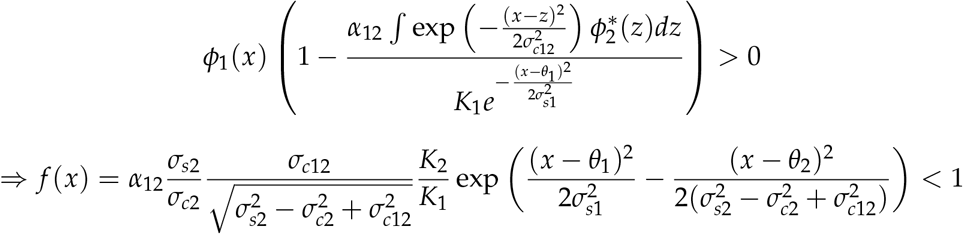

If 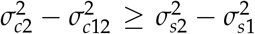 (inequality *A*), then lim_*x*→±∞_ *f*(*x*) = 0 and the inequality holds for some *x*. Else, lim_*x*→±∞_ *f*(*x*) = ∞ and the condition holds only if min *f*(*x*) < 1. This gives the invasion criteria for species 1,

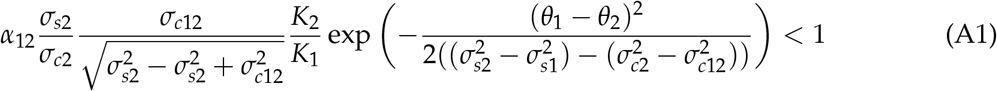

Since the system is symmetric, a similar analysis yields the invasion criteria for species 2.

## Exploiter-victim

For exploiter to coexist with the victim, the exploiter should be able to grow from low density in a population of victim. In the absence of the exploiter and strong competition among victims 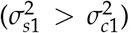, the victim trait distribution will be 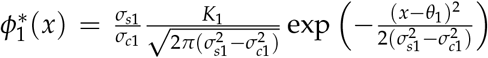. From equation (8), the condition for coexistence is,

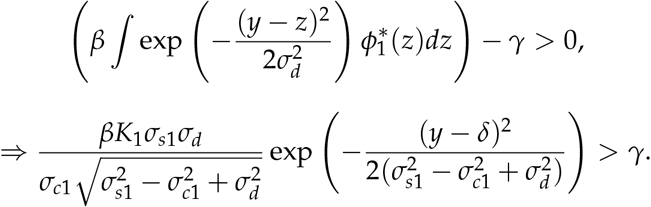

This is a Gaussian function. Therefore for some trait *y* of the exploiter to coexist with the victim, the maximum value of the function should satisfy the condition. This gives the sufficient conditions for coexistence,

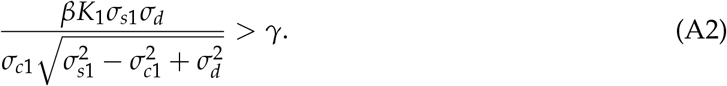

equation (A2) is the sufficient condition for coexistence when intraspecific variation is allowed to change over time, and intraspecific competition in victim is stronger than the stabilizing selection acting on them 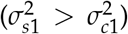. If 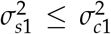, then the sufficient condition for coexistence is *β* > *γ* which is identical to the model with no intraspecific variation.

## Mutualism

To obtain conditions for the two species to exist at finite population sizes at equilibrium, we find a sequence of trait distributions for species 1 and species 2 which is the upper bound for the dynamics. The equilibrium population will be finite only if the sequence of population sizes obtained from the sequence of trait distributions converges. If *ϕ*_1_(*x*, 0) ≈ 0 and *ϕ*_2_(*y*, 0) ≈ 0, *ϕ*_1_(*x*, *τ*) will be smaller than its one species equilibrium trait distribution. Assume 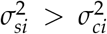. Then, for some small *τ*_1_,

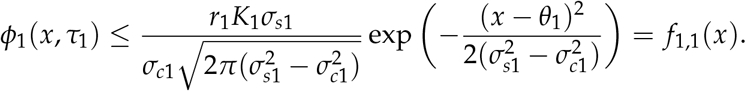

Mutualistic benefit for species 2 at time *τ* is bounded by

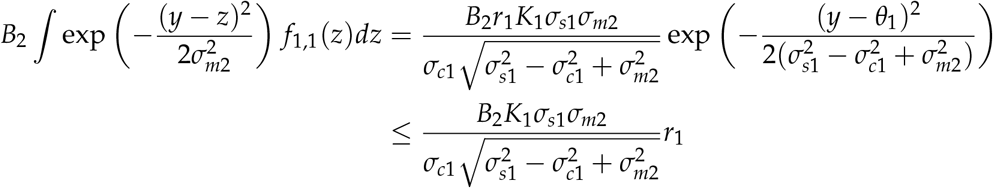

Therefore for *τ_i_* < *τ_i_*_+1_,

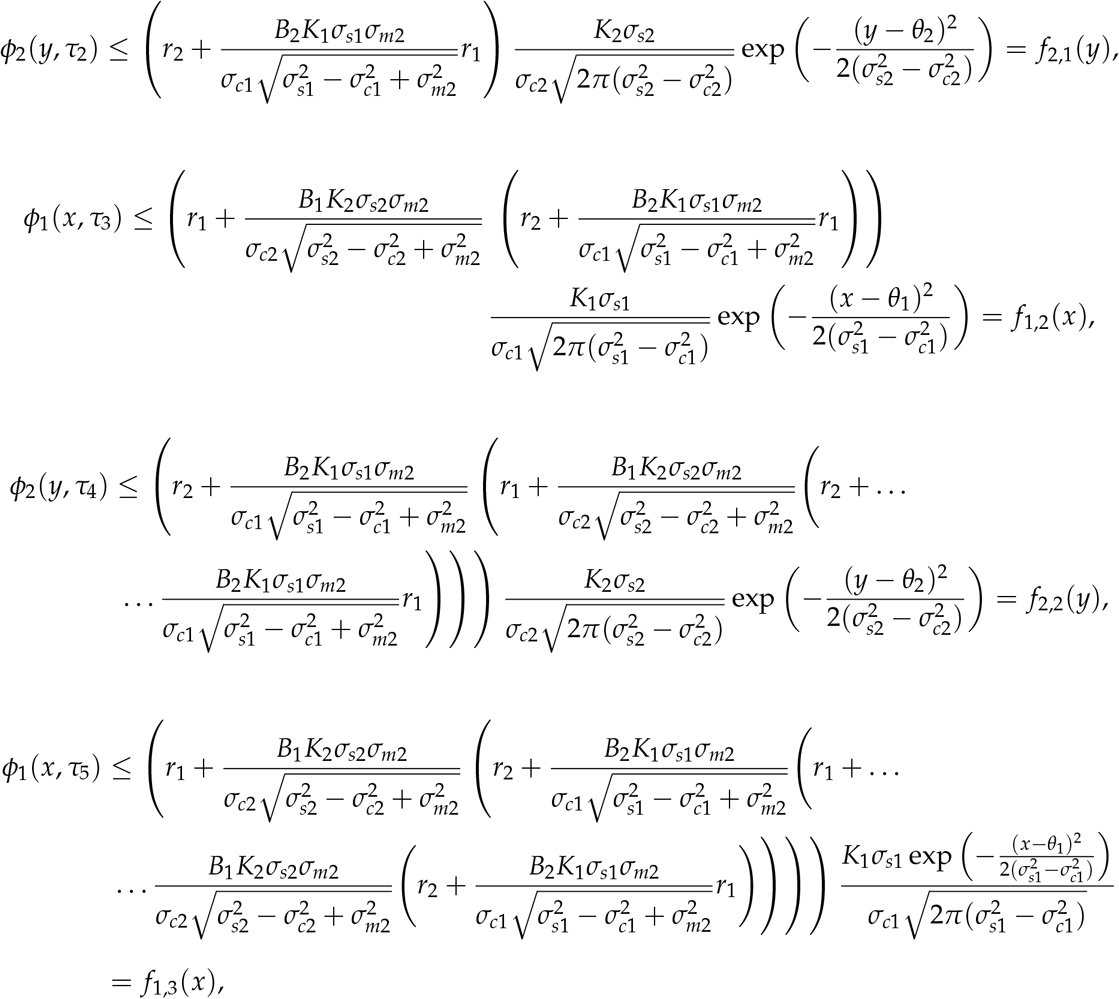

The time series of population sizes of the two species are bounded by the sequences *N*_1,*i*_ = *∫ f*_1,*i*_(*x*)*dx* and *N*_2,*i*_ = *∫ f*_2,*i*_(*y*)*dy* respectively. From the sequence of trait distributions, we can infer that

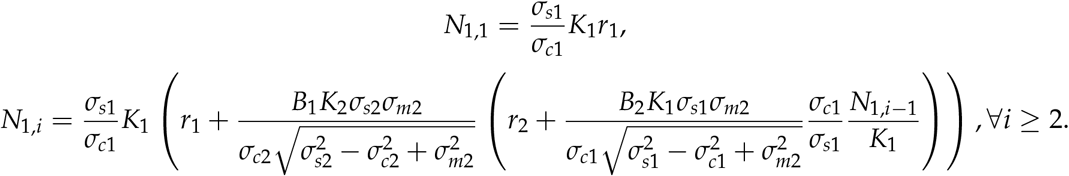

The equilibrium population size is finite only if these sequences converge. Real-valued sequences converge if and only if they are Cauchy.

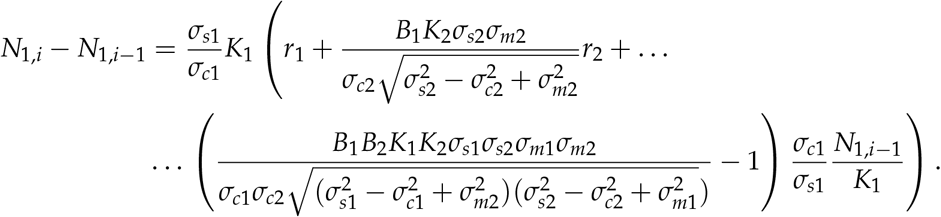

Since *N*_1,*i*_ is an increasing sequence, for it to be Cauchy,

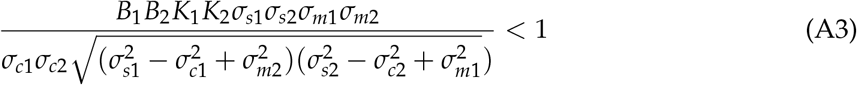

The condition for convergence of *N*_2,*i*_ is identical. Equation (A3) is a necessary condition for the equilibrium population of the two species to be finite.

## Appendix B: Numerical Method

We solved our dynamic equations using a finite difference method (Doebeli, 2011; Kreyszig *et al.*, 2011; Simmons and Krantz, 2007). Specifically, we first truncate the phenotype space to a finite interval [−*λ*, *λ*]. *λ* needs to be chosen such that *ϕ* and 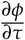 are small. Second, we partition the truncated phenotype space into intervals of length *l*. This gives a partition of size *N* = 2[*λ*/*l*]. We can now discretise the dynamic equations. For example, in the two species competition model (4),

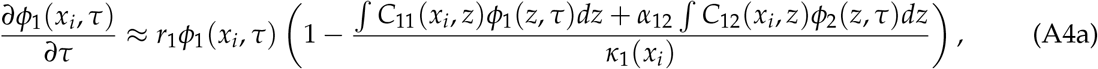

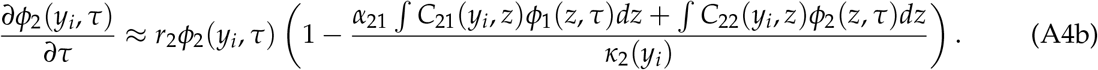

Here, the integrals are over [−*λ*, *λ*] and 1 ≤ *i* ≤ *N*. Finally, the integrals can be computed using trapezoidal rule over the same partition and the derivatives can be approximated using the Euler forward method. This leads to the iterative equations:

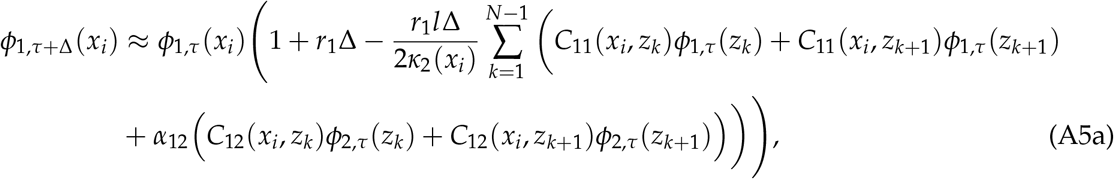

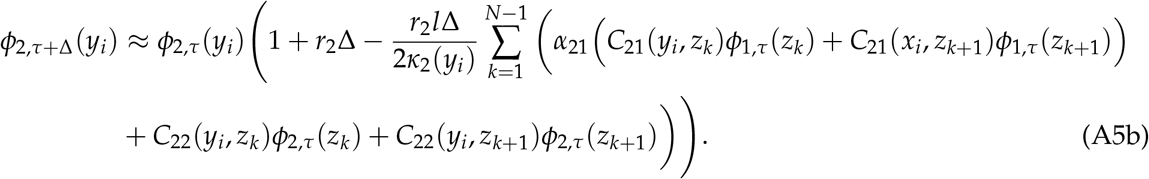

For small Δ and *l*, these equations converge. Numerical convergence can be made faster by using adaptive time steps. This is achieved by halving the time steps and reiterating one step whenever population densities becomes negative. Time steps can also be occasionally doubled if population densities remains positive over several time steps.

Setting *α*_12_ = *α*_21_ = 0 reduces the two-species competition model to two independent single-species model (equation (2)). We then confirmed that the equilibrium phenotypic distribution we obtain using the numerical method matches the analytical solution of the single-species model for different parameter choices.

This same method was applied to the exploiter-victim and mutualism models.

## Literature Cited

Abrams, P. A. (2000). The Evolution of Predator-Prey Interactions: Theory and Evidence. Annual Review of Ecology and Systematics, 31(1), 79–105.

Abrams, P. A. and Matsuda, H. (1994). The evolution of traits that determine ability in competitive contests. Evolutionary Ecology, 8(6), 667–686.

Akçay, E. (2016). Evolutionary Models of Mutualism. In Mutualism, pages 57–76. Oxford University Press.

Albert, C. H., Grassein, F., Schurr, F. M., Vieilledent, G., and Violle, C. (2011). When and how should intraspecific variability be considered in trait-based plant ecology? Perspectives in Plant Ecology, Evolution and Systematics, 13(3), 217–225.

Allen, W. J., Meyerson, L. A., Flick, A. J., and Cronin, J. T. (2018). Intraspecific variation in indirect plant–soil feedbacks influences a wetland plant invasion. Ecology, 99(6), 1430–1440.

Anderson, B. and Johnson, S. D. (2008). The geographical mosaic of coevolution in a plant-pollinator mutualism. Evolution, 62(1), 220–225.

Armstrong, R. A. and McGehee, R. (1980). Competitive Exclusion. The American Naturalist, 115(2), 151–170.

Austin, M. W. and Dunlap, A. S. (2019). Intraspecific variation in worker body size makes North American bumble bees (Bombus spp.) less susceptible to decline. The American Naturalist, page 704280.

Barabás, G., D’Andrea, R., and Stump, S. M. (2018). Chesson’s coexistence theory. Ecological Monographs, 88(3), 277–303.

Barton, N. H. (1986). The maintenance of polygenic variation through a balance between mutation and stabilizing selection. Genetical Research, 47(3), 209–216.

Benkman, C. W., Parchman, T. L., Favis, A., and Siepielski, A. M. (2003). Reciprocal selection causes a coevolutionary arms race between crossbills and lodgepole pine. American Naturalist, 162(2), 182–194.

Bolnick, D. I., Amarasekare, P., Araújo, M. S., Bürger, R., Levine, J. M., Novak, M., Rudolf, V. H. W., Schreiber, S. J., Urban, M. C., and Vasseur, D. A. (2011). Why intraspecific trait variation matters in community ecology. Trends in Ecology and Evolution, 26(4), 183–192.

Breza, L. C., Souza, L., Sanders, N. J., and Classen, A. T. (2012). Within and between population variation in plant traits predicts ecosystem functions associated with a dominant plant species. Ecology and Evolution, 2(6), 1151–1161.

Bronstein, J. L. (2015). Mutualism. Oxford University Press.

Brouat, C., Garcia, N., Andary, C., and McKey, D. (2001). Plant lock and ant key: Pairwise coevolution of an exclusion filter in an ant-plant mutualism. Proceedings of the Royal Society B: Biological Sciences, 268(1481), 2131–2141.

Brown, W. L. and Wilson, E. O. (1956). Character Displacement. Systematic Zoology, 5(2), 49.

Buckling, A. and Rainey, P. B. (2002). The role of parasites in sympatric and allopatric host diversification. Nature, 420(6915), 496–499.

Bulmer, M. G. (1971). The Effect of Selection on Genetic Variability. The American Naturalist, 105(943), 201–211.

Chesson, P. (2000). Mechanisms of Maintenance of Species Diversity. Annual Review of Ecology and Systematics, 31, 343–366.

Clark, J. S. (2010). Individuals and the Variation Needed for High Species Diversity in Forest Trees. Scientific Reports, 327(February), 1129–1132.

Clarke, B. C. (1979). The evolution of genetic diversity. Proceedings of the Royal Society of London - Biological Sciences, 205(1161), 453–474.

Clayton, D. H., Lee, P. L. M., Tompkins, D. M., and Brodie III, E. D. (1999). Reciprocal Natural Selection on Host-Parasite Phenotypes. The American Naturalist, 154(3), 261–270.

Davies, N. B. and Brooke, M. d. L. (1988). Cuckoos versus reed warblers: adaptation and counter-adaptations. Animal Behaviour, 36, 262–284.

Dayan, T. and Simberloff, D. (2005). Ecological and community-wide character displacement: the next generation. Ecology Letters, 8(8), 875–894.

Des Roches, S., Post, D. M., Turley, N. E., Bailey, J. K., Hendry, A. P., Kinnison, M. T., Schweitzer, J. A., and Palkovacs, E. P. (2017). The ecological importance of intraspecific variation. Nature Ecology & Evolution, 2(1), 57.

Dieckmann, U. and Doebeli, M. (1999). On the origin of species by sympatric speciation. Nature, 400(22), 354–357.

Diekmann, O. (2004). A beginner’s guide to adaptive dynamics. Banach Center Publications, 63, 47–86.

Doebeli, M. (1997). Genetic variation and the persistence of predator-prey interactions in the Nicholson-Bailey model. Journal of Theoretical Biology, 188(1), 109–120.

Doebeli, M. (2011). Adaptive diversification (MPB-48). Princeton University Press.

Doebeli, M. and Dieckmann, U. (2000). Evolutionary Branching and Sympatric Speciation Caused by Different Types of Ecological Interactions. The American Naturalist, 156(S4), S77–S101.

Doebeli, M. and Ispolatov, I. (2010). Complexity and diversity. Science (New York, N.Y.), 328(5977), 494–7.

Ellner, S. P., Snyder, R. E., Adler, P. B., and Hooker, G. (2019). An expanded modern coexistence theory for empirical applications. Ecology Letters, 22(1), 3–18.

Ferrière, R., Bronstein, J. L., Rinaldi, S., Law, R., and Gauduchon, M. (2002). Cheating and the evolutionary stability of mutualisms. Proceedings of the Royal Society B: Biological Sciences, 269(1493), 773–780.

Ferrière, R., Gauduchon, M., and Bronstein, J. L. (2007). Evolution and persistence of obligate mutualists and exploiters: Competition for partners and evolutionary immunization. Ecology Letters, 10(2), 115–126.

Fleischer, S. R., TerHorst, C. P., and Li, J. (2018). Pick your trade-offs wisely: Predator-prey eco-evo dynamics are qualitatively different under different trade-offs. Journal of Theoretical Biology, 456, 201–212.

Foster, K. R. and Kokko, H. (2006). Cheating can stabilize cooperation in mutualisms. Proceedings of the Royal Society B: Biological Sciences, 273(1598), 2233–2239.

Frankham, R. (1996). Relationship of Genetic Variation to Population Size in Wildlife. Conservation Biology, 10(6), 1500–1508.

Fricke, E. C., Tewksbury, J. J., and Rogers, H. S. (2019). Linking intra-specific trait variation and plant function: seed size mediates performance tradeoffs within species. Oikos, page oik.06494.

Futuyma, D. J. and Slatkin, M. (1983). Coevolution. Sinauer Associates.

Gause, G. F. and Witt, A. A. (1935). Behavior of Mixed Populations and the Problem of Natural Selection. The American Naturalist, 69(725), 596–609.

Gavrilets, S. (1997). Coevolutionary chase on exploiter-victim systems with polygenic characters. Journal of theoretical biology, 186(4), 527–534.

Gavrilets, S. (2004). Fitness Landscapes and the Origin of Species (MPB-41). Princeton University Press.

Gavrilets, S. and Hastings, A. (1994a). Dynamics of genetic variability in two-locus models of stabilizing selection. Genetics, 138(2), 519–532.

Gavrilets, S. and Hastings, A. (1994b). Maintenance of multilocus variability under strong stabilizing selection. Journal of Mathematical Biology, 32(4), 287–302.

Gavrilets, S. and Hastings, A. (1995). Dynamics of polygenic variability under stabilizing selection, recombination, and drift. Genetical Research, 65(1), 63–74.

Gavrilets, S. and Michalakis, Y. (2008). Effects of environmental heterogeneity on victim-exploiter coevolution. Evolution, 62(12), 3100–3116.

Gavrilets, S. and Waxman, D. (2002). Sympatric speciation by sexual conflict. Proceedings of the National Academy of Sciences of the United States of America, 99(16), 10533–10538.

Geritz, S. A. H., Kisdi, É., Meszéna, G., and Metz, J. A. (1998). Evolutionarily Singular Strategies and the Adaptive Growth and Branching of the Evolutionary Tree. Evolutionary Ecology, 12(1), 35–57.

Gómez, J. M., Perfectti, F., Bosch, J., and Camacho, J. P. (2009). A geographic selection mosaic in a generalized plant-pollinator-herbivore system. Ecological Monographs, 79(2), 245–263.

Gomulkiewicz, R., Thompson, J. N., Holt, R. D., Nuismer, S. L., and Hochberg, M. E. (2000). Hot Spots, Cold Spots, and the Geographic Mosaic Theory of Coevolution. The American Naturalist, 156(2), 156–174.

Hart, S. P., Schreiber, S. J., and Levine, J. M. (2016). How variation between individuals affects species coexistence. Ecology Letters, 19(8), 825–838.

Hausch, S., Vamosi, S. M., and Fox, J. W. (2018). Effects of intraspecific phenotypic variation on species coexistence. Ecology, 99(6), 1453–1462.

Jacquemyn, H., Vandepitte, K., Roldán-Ruiz, I., and Honnay, O. (2009). Rapid loss of genetic variation in a founding population of Primula elatior (Primulaceae) after colonization. Annals of Botany, 103(5), 777–783.

Janzen, D. H. (1980). When is it coevolution? Evolution, 34(3), 611–612.

Jung, V., Violle, C., Mondy, C., Hoffmann, L., and Muller, S. (2010). Intraspecific variability and trait-based community assembly. Journal of Ecology, 98(5), 1134–1140.

Keller, I. and Largiadér, C. R. (2003). Recent habitat fragmentation caused by major roads leads to reduction of gene flow and loss of genetic variability in ground beetles. Proceedings of the Royal Society B: Biological Sciences, 270(1513), 417–423.

Kisdi, É. (1999). Evolutionary branching under asymmetric competition. Journal of Theoretical Biology, 197(2), 149–162.

Kopp, M. and Gavrilets, S. (2006). Multilocus genetics and the coevolution of quantitative traits. Evolution, 60(7), 1321–1336.

Kostitzin, V. (1939). Mathematical biology. George G. Harrap And Company Ltd. London.

Kot, M. (2001). Elements of Mathematical Ecology. Cambridge University Press.

Kremer, C. T. and Klausmeier, C. A. (2017). Species packing in eco-evolutionary models of seasonally fluctuating environments. Ecology Letters, 20(9), 1158–1168.

Kreyszig, E., Kreyszig, H., and Norminton, E. J. E. J. (2011). Advanced engineering mathematics. Wiley.

Lande, R. (1975). The maintenance of genetic variability by mutation in a polygenic character with linked loci. Genetical Research, 26(3), 221–235.

Leimar, O., Sasaki, A., Doebeli, M., and Dieckmann, U. (2013). Limiting similarity, species packing, and the shape of competition kernels. Journal of Theoretical Biology, 339, 3–13.

Lichstein, J. W., Dushoff, J., Levin, S. A., and Pacala, S. W. (2007). Intraspecific variation and species coexistence. American Naturalist, 170(6), 807–818.

Lloyd-Smith, J. O., Schreiber, S. J., Kopp, P. E., and Getz, W. M. (2005). Superspreading and the impact of individual variation on disease emergence. Nature, 438(7066), 355–9.

Lotka, A. J. (1920). Analytical Note on Certain Rhythmic Relations in Organic Systems. Proceedings of the National Academy of Sciences, 6(7), 410–415.

May, R. (1981). Models for Two Interacting Populations. In Theoretical Ecology. Principles and Applications, pages 78–104. Blackwell Publishing Ltd.

McPeek, M. A. (2017). Evolutionary community ecology. Princeton University Press.

Mitrovski, P., Hoffmann, A. A., Heinze, D. A., and Weeks, A. R. (2008). Rapid loss of genetic variation in an endangered possum. Biology Letters, 4(1), 134–138.

Nijhawan, S., Mitapo, I., Pulu, J., Carbone, C., and Rowcliffe, J. M. (2019). Does polymorphism make Asiatic golden cat the most adaptable predator in Eastern Himalayas? Ecology, page e02768.

Nuismer, S. L. (2017). Introduction to coevolutionary theory. Macmillan Higher Education.

Nuismer, S. L. and Doebeli, M. (2004). Genetic correlations and the coevolutionary dynamics of three-species systems. Evolution, 58(6), 1165–1177.

Nuismer, S. L., Thompson, J. N., and Gomulkiewicz, R. (2000). Coevolutionary clines across selection mosaics. Evolution, 54(4), 1102–1115.

Nuismer, S. L., Doebeli, M., and Browning, D. (2005). The coevolutionary dynamics of antagonistic interactions mediated by quantitative traits with evolving variances. Evolution, 59(10), 2073–2082.

Nuismer, S. L., Gomulkiewicz, R., and Ridenhour, B. J. (2010). When is correlation coevolution? The American naturalist, 175(5), 525–537.

Pfennig, K. S. and Pfennig, D. W. (2009). Character displacement: ecological and reproductive responses to a common evolutionary problem.

Roughgarden, J. (1972). Evolution of Niche Width. The American Naturalist, 106(952), 683–718.

Roughgarden, J. (1976). Resource partitioning among competing species -A coevolutionary approach. Theoretical Population Biology, 9(3), 388–424.

Schreiber, S. J., Patel, S., and TerHorst, C. P. (2016). Evolution as a coexistence mechanism: does genetic architecture matter? The American Naturalist, pages 1–53.

Schulter, D. and McPhail, J. D. (1992). Ecological Character Displacement and Speciation in Sticklebacks. The American Naturalist, 140(1), 85–1098.

Simmons, G. F. and Krantz, S. G. (2007). Differential equations: theory, technique, and practice. McGraw-Hill Higher Education.

Slatkin, M. (1980). Ecological Character Displacement. Ecology, 61(1), 163–177.

Smith, P. J., Francis, R. I., and McVeagh, M. (1991). Loss of genetic diversity due to fishing pressure. Fisheries Research, 10(3-4), 309–316.

Soler, J. J., Martínez, J. G., Soler, M., and Møller, A. P. (2001). Coevolutionary interactions in a host-parasite system. Ecology Letters, 4(5), 470–476.

Start, D. (2019). Individual and Population Differences Shape Species Interactions and Natural Selection. The American Naturalist.

Start, D. and Gilbert, B. (2019). Trait variation across biological scales shapes community structure and ecosystem function. Ecology, page e02769.

Summers, K., McKeon, S., Sellars, J., Keusenkothen, M., Morris, J., Gloeckner, D., Pressley, C., Price, B., and Snow, H. (2003). Parasitic exploitation as an engine of diversity.

Taper, M. L. and Case, T. J. (1992). Models of Character Displacement and the Theoretical Robustness of Taxon Cycles. Evolution, 46(2), 317–333.

Thompson, J. N. (1994). The coevolutionary process. University of Chicago Press.

Thompson, J. N. (2005). The geographic mosaic of coevolution. University of Chicago Press.

Toju, H. and Sota, T. (2006). Adaptive divergence of scaling relationships mediates the arms race between a weevil and its host plant. Biology Letters, 2(4), 539–542.

Turchin, P. (2003). Complex Population Dynamics: A Theoretical/Empirical Synthesis (MPB-35). Princeton University Press.

Vellend, M. (2006). The consequences of genetic diversity in competitive communities. Ecology, 87(2), 304–311.

Violle, C., Enquist, B. J., McGill, B. J., Jiang, L., Albert, C. H., Hulshof, C., Jung, V., and Messier, J. (2012). The return of the variance: Intraspecific variability in community ecology.

Walsh, B. and Lynch, M. (2018). Maintenance of Quantitative Genetic Variation. In Evolution and Selection of Quantitative Traits, volume 1. Oxford University Press.

Waxman, D. and Gavrilets, S. (2005). 20 Questions on Adaptive Dynamics. Journal of Evolutionary Biology, 18(5), 1139–1154.

Winkelmann, K., Genner, M. J., Takahashi, T., and Rüber, L. (2014). Competition-driven speciation in cichlid fish. Nature Communications, 5(1), 3.

Yoder, J. B. and Nuismer, S. L. (2010). When Does Coevolution Promote Diversification? The American naturalist, 176(6), 802–817.

Zu, J. and Wang, J. (2013). Adaptive evolution of attack ability promotes the evolutionary branching of predator species. Theoretical Population Biology, 89, 12–23.

